# Sample size calculation for a NanoString GeoMx spatial transcriptomics experiment to study predictors of fibrosis progression in non-alcoholic fatty liver disease

**DOI:** 10.1101/2023.01.20.524846

**Authors:** Maria Ryaboshapkina, Vian Azzu

**Affiliations:** Translational Science and Experimental Medicine, Research and Early Development, Cardiovascular, Renal and Metabolism (CVRM), BioPharmaceuticals R&D, AstraZeneca, Gothenburg, Sweden; Translational Science and Experimental Medicine, Research and Early Development, Cardiovascular, Renal and Metabolism (CVRM), BioPharmaceuticals R&D, AstraZeneca, Cambridge, UK

**Keywords:** Spatial transcriptomics, NanoString GeoMx Whole Transcriptome Atlas, sample size calculation, non-alcoholic fatty liver disease

## Abstract

Sample size calculation for spatial transcriptomics is a novel and understudied research topic. Prior publications focused on powering spatial transcriptomics studies to detect specific cell populations or spatially variable expression patterns on tissue slides. However, power calculations for translational or clinical studies often relate to the difference between patient groups, and this is poorly described in the literature. Here, we present a stepwise process for sample size calculation to identify predictors of fibrosis progression in non-alcoholic fatty liver disease as a case study. We illustrate how to infer study hypothesis from prior bulk RNA-sequencing data, gather input requirements and perform a simulation study to estimate required sample size to evaluate gene expression differences between patients with stable fibrosis and fibrosis progressors with NanoString GeoMx Whole Transcriptome Atlas assay.

**Key points:** Spatial transcriptomics is predominantly used in unpowered exploratory research.

We demonstrate that spatial transcriptomics studies with NanoString GeoMx Whole Transcriptome Atlas can be powered to address differences between patient groups and illustrate the sample size calculation using fibrosis progression in non-alcoholic fatty liver disease as an example.

We note that established software implementing negative binomial mixed effect models lme4 and GLMMadaptive yield more similar fold change estimates to simulated data than the recently published software GeoDiff.

## Introduction

Spatial transcriptomics offers significant advantages over bulk or single cell transcriptomics. Bulk transcriptomics is unable to reveal tissue-wide patterning and disease-promoting cell niches. Single cell omics typically requires fresh tissue, whose dissociation can lead to significant cell type biases. Both these factors can be overcome with spatial transcriptomics, which additionally offers the ability to use widely available formalin-fixed paraffin-embedded (FFPE) tissue.

Two spatial RNA sequencing technologies are commercially available: Visium spatial gene expression from 10X Genomics[1] and GeoMx Digital Spatial Profiler from NanoString [2, 3].

The Visium platform uses functionalized slides with printed barcoded oligo capture probes onto which RNA from permeabilised tissue can bind. This can subsequently be sequenced using a polyA approach. Visium is limited by pre-determined sequenceable spot sizes of 55μm with 100μm spacing between spot centres.

The GeoMX platform extends more flexibility in enabling the operator to select regions of interest/illumination (ROI) from single cell analysis to whole regions based on histology or protein probes. This platform links complementary sequence probes to a unique barcode through a UV-cleavable linker. Hybridised sequence probes on the tissue of interest can then be released from tissue sections through the application of UV only to selected ROIs. The GeoMX platform allows addition of custom targets of interest such as transcript variants. GeoMx Human Whole Transcriptome Atlas enables targeted sequencing of 18,000 genes [3]. While both Visium and NanoString GeoMx are compatible with FFPE tissue, sequencing efficiency with Visium is reduced in FFPE compared to fresh frozen tissue while no such limitation is reported for NanoString GeoMx [4].

Sample size calculation is a crucial step in planning spatial transcriptomics studies. Yet, limited methodological research is available on this topic, especially in clinical specimens. Previous studies reporting sample size calculation for spatial transcriptomics focused on detection or co-localization of specific cell populations on the slide or identification of spatially variable features [5–7]. Specifically, despite the widespread use of the technology in unpowered exploratory research (https://nanostring.com/resources/publications/), only a handful of studies with NanoString GeoMx report dedicated sample size calculation based on pilot experiments but do not outline the requirements and statistical considerations in detail [8, 9].

Here, we illustrate how to gather required inputs (formulate study hypothesis and estimate effect size based on historic data) and perform sample size estimation for comparison between patient groups with NanoSting GeoMx Whole Transcriptome Atlas using fibrosis progression in non-alcoholic fatty liver disease as a case study.

Non-alcoholic fatty liver disease (NAFLD) affects up to a third of the population [10] and covers a spectrum of pathology from simple steatosis (NAFL) to steatohepatitis (NASH) and advanced fibrosis [11]. Progressed NAFLD places patients at risk of liver cancer and is strongly associated with the metabolic syndrome (MetS, comprising obesity, insulin resistance, type 2 diabetes mellitus, hypertension) and coronary heart disease [12].

NAFLD has been shown to progress to fibrosis both in patients with NAFL and NASH [13, 14], with fibrosing disease being the leading predictor of poor outcomes/survival [15, 16]. To date, no studies have been able to show clinical, biochemical or histological predictors of fibrosis progression in cohorts of with serial liver sampling of the same individual [13, 14, 16–19]. A combination of followup biopsy analysis and cross-sectional studies indicate that weight gain [13], metabolic co-morbidities such as diabetes [13, 14, 17, 19]and at-risk genotypes, for example in PNPLA3, TM6SF2, MBOAT7 [20, 21] place patients at higher risk of NAFLD fibrosis progression. Non-genetic molecular mechanisms placing the patient at risk of fibrosis progression are particularly poorly elucidated and limited to hepatic gene expression-based or serum protein-based risk signatures also found in HCV fibrosis progression [22]. Thus, further research is required to understand predictors fibrosis progression in NAFLD and, ultimately, adjust clinical management of higher risk patients.

## Methods

### Initial hypothesis generation

First, we performed analysis to identify which spatial expression patterns could serve as the primary and secondary endpoints.

We explored bulk liver RNA-sequencing cohort from Fujiwara *et al*. (GSE193066 [23]). Fibrosis progressors and fibrosis regressors had at least +1 and −1 fibrosis stage in their circa 2 year follow-up biopsies. The cohort included 28 individuals with stable fibrosis stage, 15 fibrosis regressors and 15 fibrosis progressors. We sought to identify an association between a higher-level gene expression pattern and some element of liver microanatomy or histology. Therefore, we did not make any attempt to control false discovery rate at the level of individual genes or pathways in this analysis. We also used the published DESeq2 rlog-transformed data set as-is without re-processing. However, we did check that results obtained with linear regression on rlog transformed data and on raw counts with DESeq2 [24] were overall consistent, especially with respect to log2 fold change estimates, with this sample size in another cohort GSE135251 [25] where both types of data were readily available (**Figure S1**). Candidate genes were identified with linear regression on rlog data adjusted for baseline fibrosis stage in GSE193066. The genes were required to have opposite direction of log2 fold change in fibrosis progressors and fibrosis regressors compared to stable fibrosis individuals and raw p < 0.05 in both comparisons. The candidate genes were semi-manually grouped into broad marker sets using AmiGO 2 [26] and PANTHER [27] based on GO [28, 29] and Reactome [30] pathway definitions and hierarchical relationships between the ontology terms. We calculated a summary score for each of these broad candidate marker sets with Kuppe’s method [31]. We verified that the expression of each candidate marker set as a whole, differed between fibrosis progressors, stable fibrosis and fibrosis regressors using linear regression adjusted for both baseline fibrosis stage and baseline NAS score.

The candidate marker sets were derived from bulk liver. As the next step, we traced potential sources of these expression signals within the liver. We calculated summary scores for liver cell types based on markers from GSE136103 [32], hepatocyte zones in NASH liver [33] and for extracellular matrix (ECM, “core matrisome” markers from [31] in the Fujiwara *et al*. cohort (GSE193066 [23]) and in published Visium liver spatial transcriptomics samples [34–36]. Visium data was SCT-transformed prior to analysis. We assumed “guilt-by-association”: if summary scores correlated both in bulk and in spatial transcriptomics samples (Spearman method), then the expression signals co-localized.

### Simulation study

Non-tumour liver samples adjacent to hepatocellular carcinoma [34] had well-defined fibrosis areas and hepatocyte fraction and satisfied the 100 reads/μm^2^ recommendation by NanoString. We used these five samples for the simulation study.

We defined fibrotic niche as Visium spots that were in the 85 percentile or above for ECM score and at or below the 15 percentile of the hepatocyte score. Similarly, hepatocyte spots were at or above the 20 percentile for hepatocyte score and at or below the 80 percentile of the ECM score. The remaining spots were enriched in other cell types (e.g., cholangiocytes) or had low cell density. This resulted in 2.6-7.1% fibrotic niche, which was within the NAFLD fibrosis range [37], and 63-68% hepatocyte area, which corresponded to the lower range of literature estimates for the hepatocyte fraction by liver volume [38–40]. In total, we obtained 819 fibrotic niche spots and 12,074 hepatocyte spots across the 5 samples. To verify that our approach to tissue segmentation was reasonable, we conducted differential expression analysis between “bulk” tissue constructed by summing up counts from fibrotic niche and hepatocyte spots per patient. As expected, fibrotic niche was enriched in markers of diverse non-parenchymal cell types (mesenchymal, immune etc) and hepatocyte fraction was enriched in hepatocyte markers (**Data S1**).

We translated the expression changes in summary scores from bulk tissue into expression changes within fibrotic niche and hepatocyte fraction. We found two representative markers PON1 and FLNA that had the most similar spatial expression pattern to the two summary scores identified in initial analysis. All subsequent calculations were based on these two genes.

We spiked-in increasingly large fractions of read counts to PON1 in hepatocytes and FLNA in the fibrotic niche. For example, spiked-in fraction r0.1 for FLNA corresponded to FLNA read counts in each fibrotic niche spot in sample i + 0.1*median(FLNA read counts across fibrotic niche spots in sample i), where i =1…5. We summed up read counts across all spots per sample to obtain “bulk” tissue. We calculated log2 fold change between spiked-in “bulk” and unmodified “bulk” tissue with DESeq2 using the patient as a fixed effect covariate and constructed calibration curves. We found the point on each calibration curve where fold change in “bulk” tissue matched log2 fold change observed in the initial hypothesis-generating analysis in the Fujiwara *et al*. cohort [23]. This determined the magnitude of spike-in that we used in the simulation study.

We took the unmodified 12,074 hepatocyte spots and 819 fibrotic niche spots across the five Visium samples as our spot “population” in “stable fibrosis” individuals. The same spots but spiked-in with fractional read counts r0.43 for PON1 and r2.9 for FLNA, respectively, were our spot “population” in “fibrosis regressors” and “fibrosis progressors”, respectively.

Initial Visium spot diameter of 55μm contains 8 to 20 cells [34] and is not recommended by NanoString because it does not support reliable differential expression (“The GeoMx® Human Whole Transcriptome Atlas for the Digital Spatial Profiler: Design, Performance, and Experimental Guidelines” document from www.nanostring.com/GeoMxDSP). Fibrotic niche was the smaller tissue fraction in our experimental plan and it determined the range of diameters that we evaluated. The width of fibrotic niche in the refence data rarely exceeded 2-3 Visium spots (55μm*2 + 45 μm distance between spots = 155μm, 55μm*3 + 2*45μm = 255μm). Reference estimates of fibrotic niche size reported in NAFLD were in the <200μm range [37, 41, 42]. Based on this prior data, we considered 165μm as maximum spot diameter. The amount of input material (cells) is determined by tissue area that is subjected to sequencing. The relationship between spot diameter and area is quadratic, so we aggregated 2 Visium spots to mimic 80μm diameter, 4 Visium spots to mimic 110|Lim, and 9 Visium spots to mimic 165 μm.

In NanoString terminology, a spot is called “region of illumination”, or ROI, and corresponds to the small piece of tissue from which barcodes are harvested for sequencing. We use the term “ROI” to describe the synthetic larger-diameter NanoString data and the term “spot” to describe original 55μm Visium spots.

We drew random subsamples from our unmodified and spiked-in Visium spot “populations” and summed up counts from neighbouring spots to mimic NanoString ROIs with larger diameter. We made 100 synthetic data sets for each condition (endpoint x N patients x N ROIs x ROI diameter). In the original Visium data, correlations between global gene expression profiles in spots within one patient were not stronger on average than between spots from different patients. So, the new synthetic larger ROIs were randomly assigned to new “patient” or a new ROI from an existing “patient”. The simulated data sets were restricted to genes measured on the Nanostring GeoMx Human Whole Transcriptome Atlas assay (GPL32201). Genes that were part of the assay and had median expression <0.05 TPM in all GTEx tissues (https://gtexportal.org/home/ [43]) were used as negative control “background probes” for the GeoDiff method [44].

Since spatial transcriptomics is a new technology and the NanoString software GeoDiff [44] has not been extensively validated yet, we wanted to increase confidence in the results and applied three methods to test differences in expression levels of FLNA and PON1 between spiked-in and unmodified data sets. Conceptually, all three methods relied on negative binomial mixed effect regression models with background or effective library size treated as a threshold or an offset. Therefore, expression of a gene of interest was corrected for background. In all cases, patient group was treated as the fixed effect (e.g. “fibrosis progressor” vs “stable fibrosis”) and patient as the random intercept. Effective library size was calculated with edgeR [45].

#### Method 1, GeoDiff [44]

GeoDiff::fitNBthmDE with gene count ~ group + (1|patient) after following the processing steps including the background estimation as described in the package vignette (https://bioconductor.org/packages/release/bioc/vignettes/GeoDiff/inst/doc/Workflow_WTA_kidney.html).

#### Method2, glmer.nb [46]

lme4::glmer.nb with gene count ~ offset(natural logarithm of effective library size) + group + (1|patient)

#### Method 3, GLMMadaptive LRT [47, 48]

Likelihood ratio test (LRT) between full model assuming that there is a group difference and the null model assuming that gene expression is explained by technical and within-patient variability.

Full model: gene count ~ offset(natural logarithm of effective library size) + group + (1|patient)

Null model: gene count ~ offset(natural logarithm of effective library size) + (1|patient)

We summarized our simulation setup in **Table 1**. For the final simulation run, the numbers of repeats/random data sets per condition was increased to 1,000.

**Table 1.**
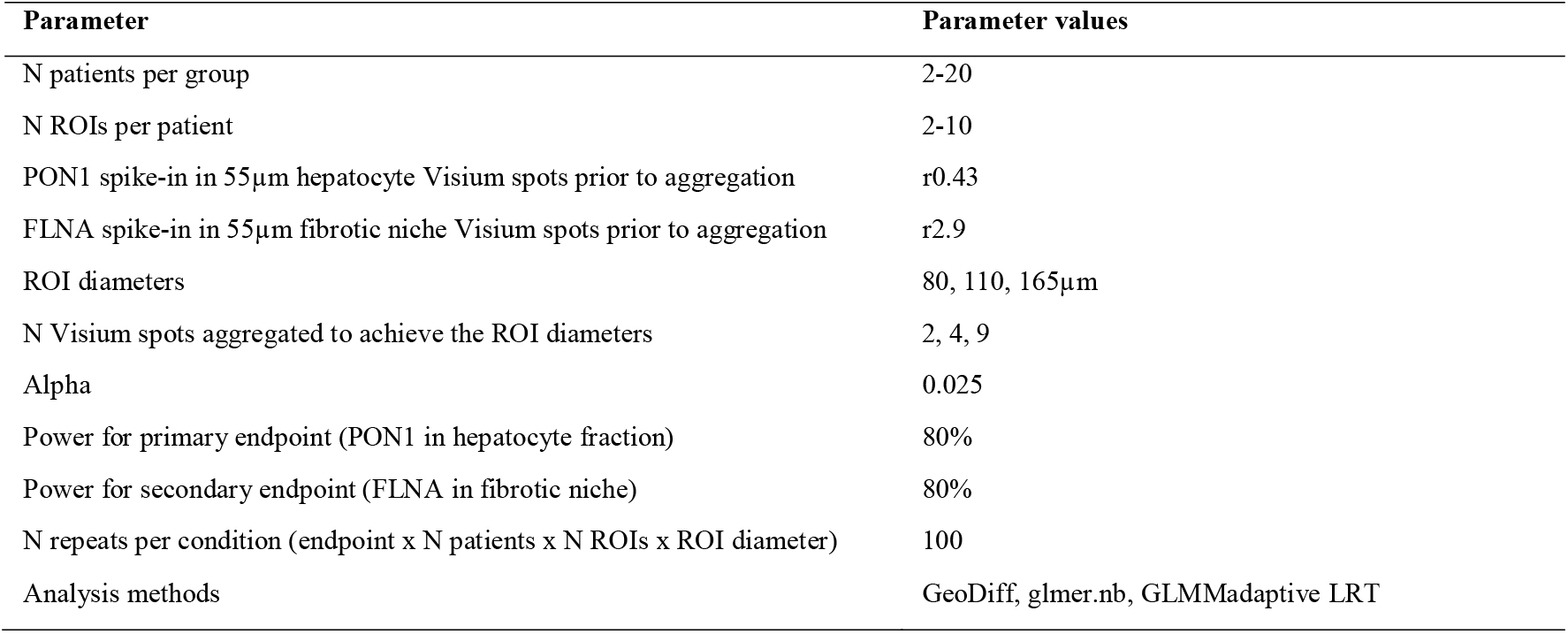
Setup of the initial simulation.

### Software

All analyses were conducted in R 4.1.0 [49], Bioconductor version 3.14. Research scripts are enclosed with this article (**Data S2**). Spatial transcriptomics data was handled with Seurat v4.1.1 [50]. Figures were made with ggplot2 [51] and viridis [52].

### Data availability

Bulk RNA-sequencing GSE193066 [23] and spatial transcriptomics [34–36] were publicly available.

## Results

### Study endpoints

Sample size calculation requires definition of the study endpoint. We used a combination of historic bulk RNA-sequencing (GSE193066 [23]) and spatial transcriptomics data [34–36] to identify spatial gene expression pattens in baseline biopsies that correlated with fibrosis outcome in NAFLD.

At baseline, 108 genes had higher expression in fibrosis regressors and lower expression in fibrosis progressors compared to stable fibrosis individuals. Among these genes, 44 (40.7%) were associated with different aspects of lipid metabolism. By contrast, 187 genes had higher baseline expression in fibrosis progressors and lower baseline expression in fibrosis regressors. Among these genes, 89 (47.6%) were associated with actin cytoskeleton, cell projections such as cell leading edge and RHO GTPase activity. We used umbrella terms “lipid metabolism” and “cytoskeleton” to denote these 44 and 89 genes (**Data S3**).

The identified gene sets were aggregated into summary scores. Analysis of the summary scores confirmed that fibrosis progressors had lower baseline expression of lipid metabolism markers (**Figure 1A**) and higher expression of cytoskeleton markers (**Figure 1B**) irrespective of their disease severity at baseline.

**Figure 1.**
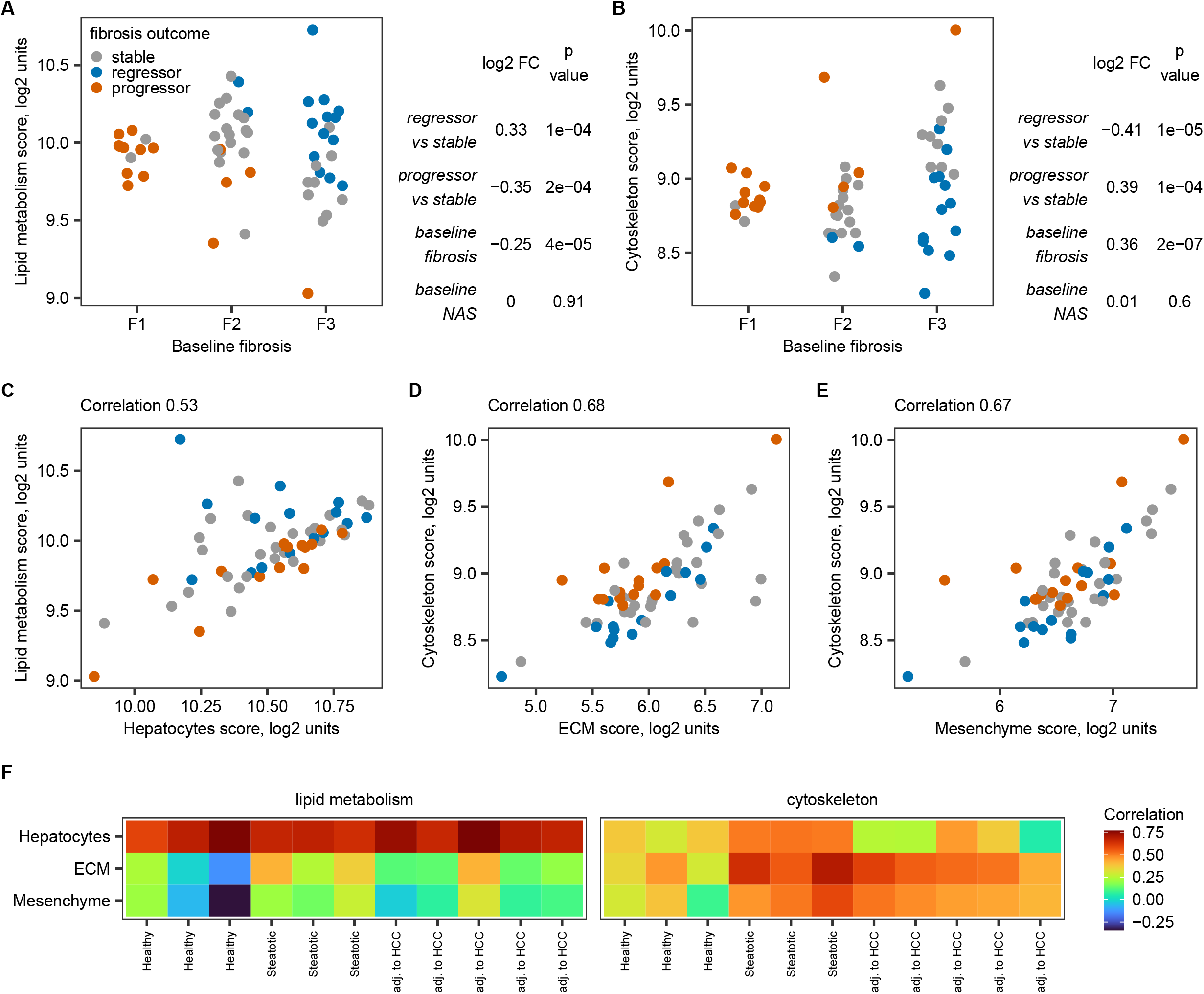
Initial hypothesis generation. (A) Relative expression of lipid metabolism score in baseline biopsies stratified by 2-year outcome and baseline fibrosis stage in bulk NAFLD liver GSE193066. N stable-fibrosis individuals = 28, fibrosis progressors = 15 and fibrosis regressors = 15. Table on the side shows coefficients from the multivariate linear regression model, in which relative expression score is a function of 2-year outcome, fibrosis stage and NAS score at baseline. (B) Relative expression of cytoskeleton score in baseline biopsies stratified by 2-year outcome and baseline fibrosis stage in bulk NAFLD liver GSE193066. Notations are as in panel (A). Possible source of expression signal (C-E) in NAFLD bulk liver GSE193066 and (F) in spatial transcriptomics experiments. Correlations in spatial transcriptomics were computed across the total tissue area within each sample separately.

Lipid metabolism score correlated with hepatocyte score (**Figure 1C**) while cytoskeleton score correlated with extracellular matrix (ECM) (**Figure 1D**) and mesenchyme scores (**Figure 1E**) in bulk NAFLD livers. Lipid metabolism score co-localized with the hepatocyte fraction in spatial transcriptomics on both normal and diseased livers. By contrast, cytoskeleton score co-localized with fibrosis areas marked by high expression of ECM and mesenchyme scores in diseased livers (**Figure 1F**). We could not further deconvolute which cell type of the mesenchymal lineage (stellate cells, activated stellate cells, vascular smooth muscle cells or fibroblasts) could lead to co-localization of this expression signal.

Based on this initial analysis, we formulated two study hypotheses:

1. Fibrosis regressors have higher baseline expression of the lipid metabolism score in hepatocytes than stable-fibrosis individuals.
2. Fibrosis progressors have higher baseline expression of the cytoskeleton score in the fibrotic niche than the fibrosis-stable patients

To facilitate reading, the study hypotheses were formulated relative to the patient group with higher expression. However, the differences in lipid metabolism and cytoskeleton scores were observed in both directions (i.e., two sided test, 1. – higher in fibrosis regressors/lower in progressors, 2. – higher in progressors/lower in regressors) and had approximately same magnitude compared to the fibrosis-stable group. The corresponding null hypotheses were that there were no group differences.

Hypothesis 1 was prioritized as the primary endpoint because we considered isolation of fibrotic niche more technically challenging and more likely to fail. However, both hypotheses were biologically plausible [53–55], and sample sizes were calculated to address both hypotheses.

Next, we needed to turn these hypotheses into quantifiable study endpoints. Hepatocytes are the most abundant fraction of liver tissue even in individuals with advanced fibrosis [38–40], so, effect size within the hepatocyte fraction would be expected to be only slightly different from the effect size observed in bulk liver. By contrast, fibrosis areas constitute 2.8% (2.1– 3.6%) in F1, 4.3% (3.3 - 5.4%) in F2, 4.8% (3.7 −7.4%) in F3 and 12.3% (8.4 −18.5%) in F4 NAFLD fibrosis stages [37]. Thus, expression signal originating within the fibrotic niche may be “diluted” in bulk liver, and we needed to translate the effect sizes between bulk liver and fibrotic niche and hepatocyte fraction. Individual genes that contributed to the summary scores did not always match the spatial expression pattern for the whole score. We found two representative markers PON1 and FLNA that had the most similar spatial expression pattern to the lipid metabolism score and cytoskeleton score, respectively (**Figure 2**) and based the subsequent calculations on these two genes. Based on the calibration curve analysis (**Figure S2**), we formulated the primary and secondary endpoints as:

1. Fibrosis regressors have 0.42±0.02 log2 fold higher baseline expression of PON1 in hepatocytes than stable fibrosis individuals.
2. Fibrosis progressors have 1.5±0.11 log2 fold higher baseline expression of FLNA in the fibrotic niche than stable fibrosis individuals.

**Figure 2.**
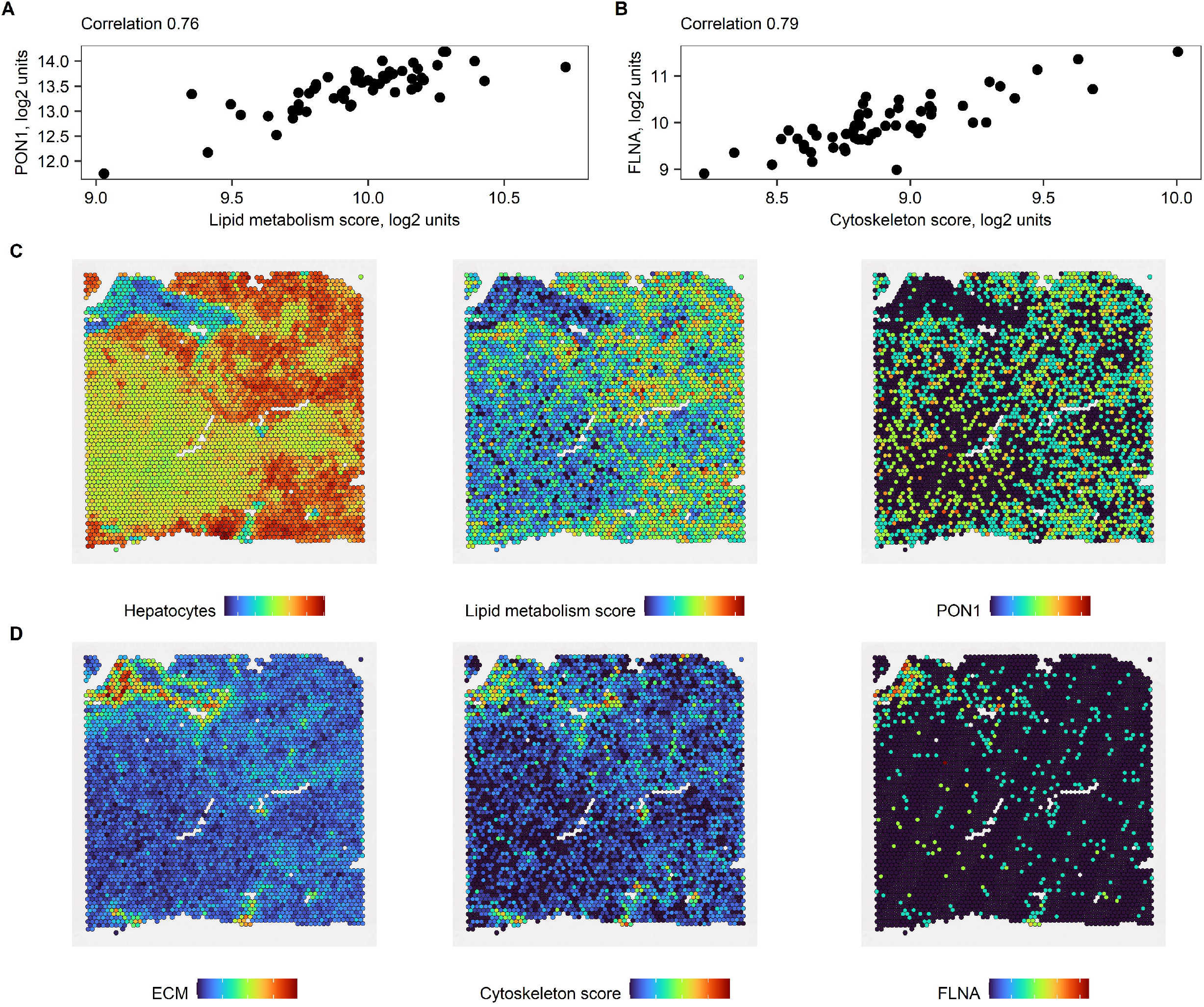
PON1 and FLNA as representative markers. Correlation between (A) PON1 and hepatocyte score and (B) FLNA and cytoskeleton score in bulk NAFLD liver GSE193066. Co-localization of (C) PON1 with lipid metabolism score in hepatocyte fraction and (D) FLNA with cytoskeleton score in fibrotic niche in representative non-tumour liver region adjacent to hepatocellular carcinoma. Colors indicate relative expression of summary marker scores. These scores are in arbitrary log2 units: blue – low, green – moderate, red – high.

### Initial simulation study

Using the NanoString GeoMx® Human Whole Transcriptome Atlas Digital Spatial Profiler, gene expression data is typically collected from 6-12 regions of illumination (ROIs) per tissue sample (MAN-10108-01_GeoMx_DSP_Experimental_Design_Guideline.pdf from www.nanostring.com/GeoMxDSP). Historic data indicated that tissue samples may have uneven thickness and cell density may be variable within tissue sample. So, the expression measurements were expected to have large technical variability in addition to biological variability of expression for PON1 in hepatocytes and FLNA in the fibrotic niche.

Given that the outcome of the experiment may be affected by the sampling variability within the tissue sample, we conducted a simulation study to accurately mirror this sampling variability and estimate the required number of patients, ROI diameter and number of ROIs per patient.

Interestingly, the three methods applied head-to-head on the same synthetic data resulted in distinct power surfaces (**Figure 3** and **4**). However, in all cases, power to detect expression differences between the groups increased with increasing number of patients per group but not necessarily with increasing number of ROIs per patient (visible in Figure 3 and 4 in each column corresponding to a fixed N patients).

**Figure 3.**
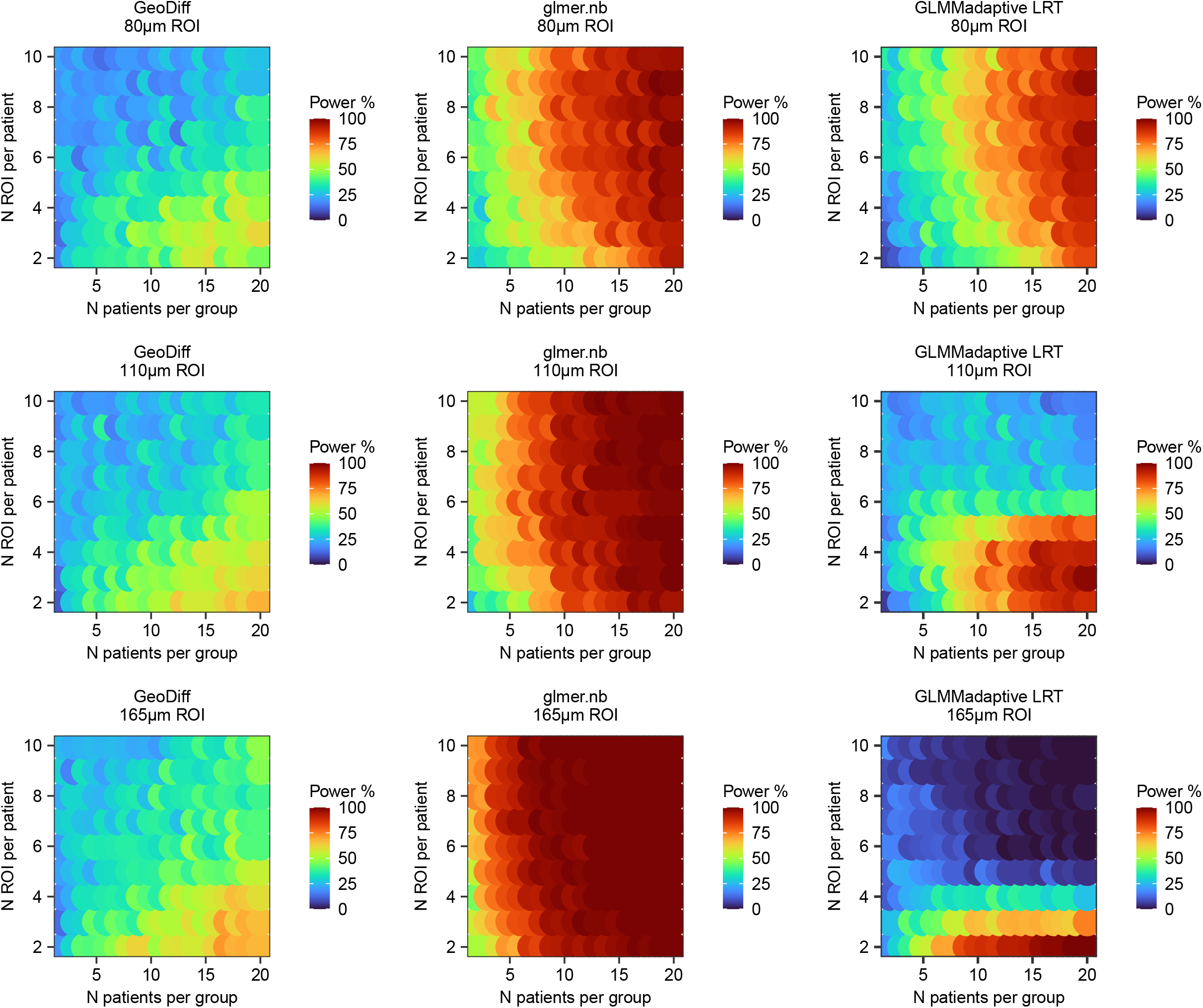
Power surface obtained for the primary endpoint (PON1 in hepatocytes) in the initial simulation. This analysis was used to narrow down the range of experimental conditions (N patients x N ROIs per patient x ROI size).

**Figure 4.**
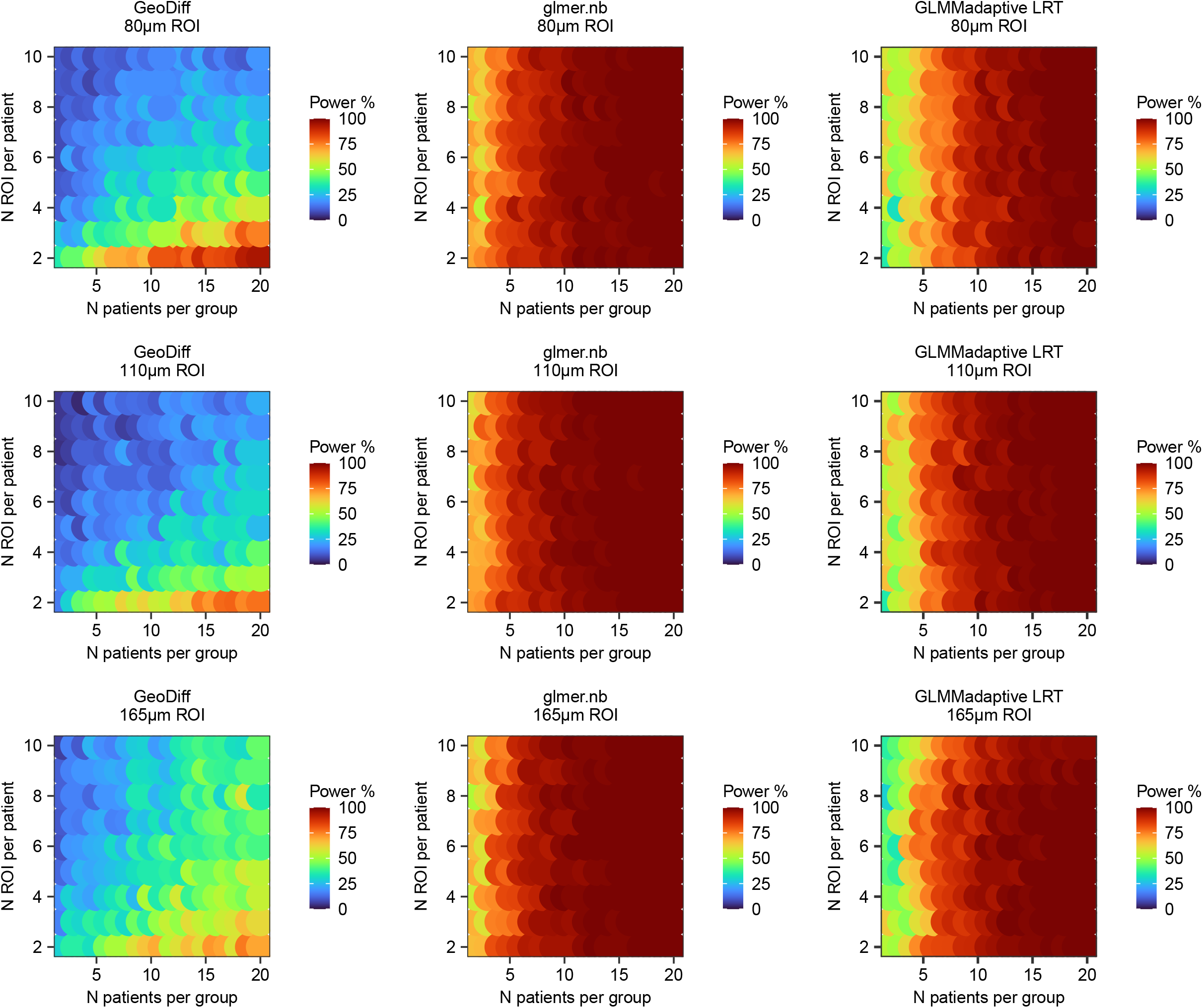
Power surface obtained for the secondary endpoint (FLNA in fibrotic niche) in the initial simulation. This analysis was used to narrow down the range of experimental conditions (N patients x N ROIs per patient x ROI size).

### Final simulation study

The initial simulation study helped us to narrow down the range of experimental conditions to the largest ROI size 165μm, two ROIs per patient and at least 4 patients per group. We repeated the simulation study but with 1,000 synthetic data sets per condition (**Table 2)**. All three methods were run on the same synthetic data, i.e., PON1 and FLNA read counts used to evaluate GeoDiff and glmer.nb and GLMMadaptive LRT methods were exactly the same. GeoDiff overestimated log2 fold change for the primary endpoint and underestimated log2 fold change for the secondary endpoint relative to the actual spike-in. Glmer.nb and GLMMadaptive LRT resulted in smaller log2 fold change estimates in the synthetic NanoString data sets than the actual spike-in but were close for the primary endpoint (PON1 in hepatocyte fraction median log2 FC 0.33 in simulation vs 0.42 in calibration curve analysis). Glmer.nb and GLMMadaptive LRT were faster than GeoDiff (e.g., final simulation for the secondary endpoint GeoDff 16.4 h, glmer.nb 6.6h and GLMMadaptive LRT 2.4h). However, we did not optimize the code, so runtimes could be reduced further. Convergence rate was 100% for glmer.nb and GeoDiff. GLMMadaptive LRT converged in 732 to 979 out of 1,000 iterations with higher failure rate on fewer patients.

**Table 2.**
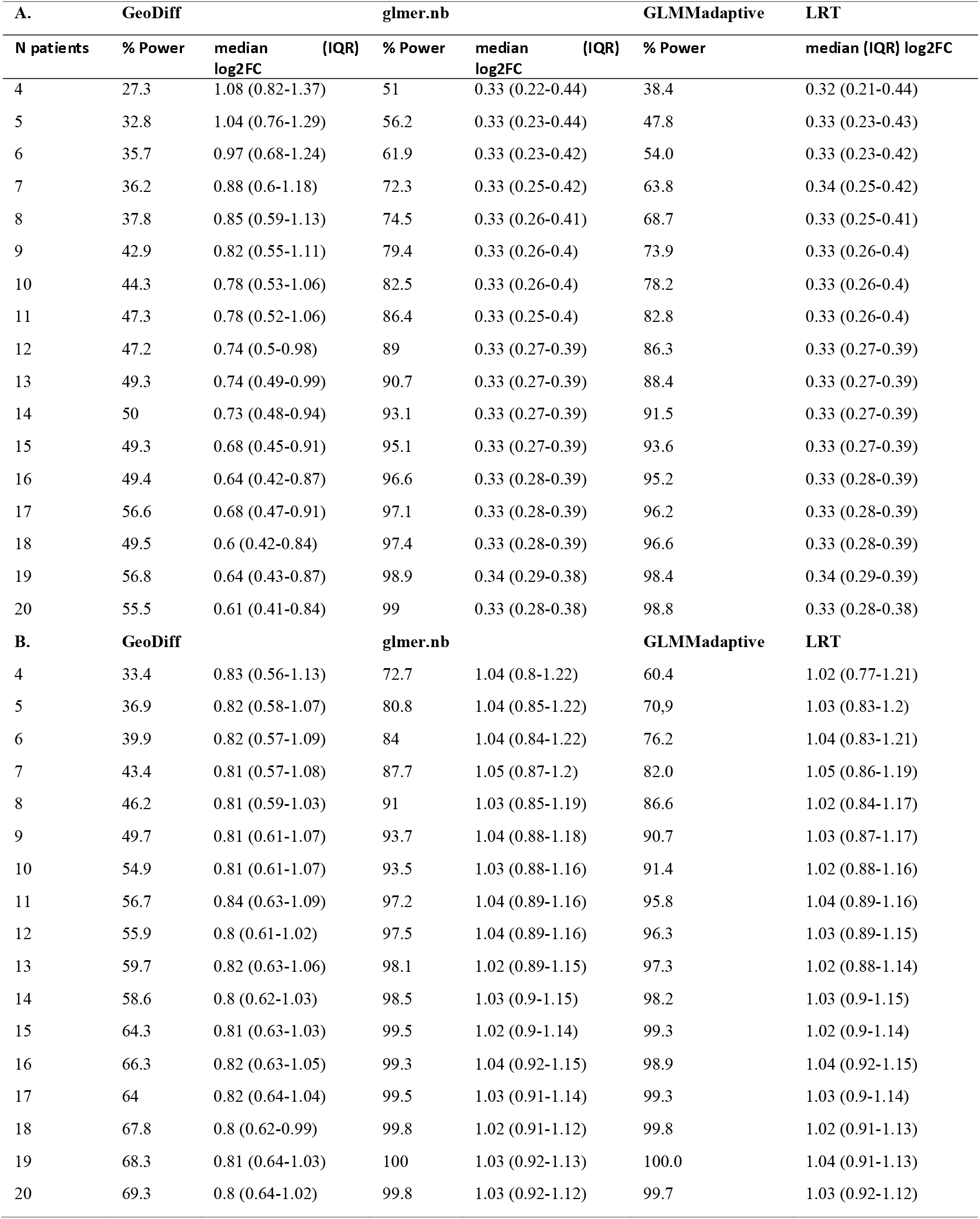
Results of the final simulation. (A) Primary endpoint PON1 in 165μm hepatocyte ROIs and 2 ROIs per patient. (B) Secondary endpoint FLNA in 165μm in fibrotic niche and 2 ROIs per patient. Number of patients is indicated per group. Log2 fold change is indicated between spiked-in and unmodified simulated data across 1,000 iterations per condition.

Taking all these factors into consideration, the final sample size was based on the glmer.nb method. We calculated that we would need (10 patients x 2 hepatocyte ROIs + 5 patients (subset of the ten) x 2 fibrotic niche ROIs) x 3 experimental groups (stable fibrosis, fibrosis regressors, fibrosis progressors) = 90 samples subjected to sequencing at baseline. If paired follow-up biopsies are profiled for exploratory analysis from the same patients and for the same tissue areas, then the total sample size for baseline and equal number of follow-up samples is 180 samples.

### Sensitivity analysis

The summary scores calculated from normalized data could also be approximated as sum of raw gene counts for all genes constituting the score. For example, Spearman correlation between lipid metabolism score calculated on normalized counts and sum of raw counts for lipid metabolism genes was median 0.79 (IQR 0.72 - 0.82) in the spatial transcriptomics samples. Sensitivity analysis using sum of gene counts resulted in similar fold change estimates in calibration curve analysis (**Figure S3**) and similar sample size estimates (**Table S1**). Fourteen and four patients per group were sufficient to reach 80% power for testing group differences in expression of the lipid metabolism and cytoskeleton genes in the hepatocyte fraction and fibrotic niche, respectively.

## Discussion

Sample size calculation is a crucial step in planning studies that are appropriately powered to answer the research question. Limited methodological research is available for sample size calculation for spatial transcriptomics in clinical research. In this study, we outline how to gather required inputs (e.g. estimate effect size based on historic data) and perform sample size estimation for comparison between patient groups. Our biological question of interest was the difference in hepatic gene expression levels between patient groups (fibrosis progressors, fibrosis regressors and stable fibrosis patients) assayed with NanoString GeoMx spatial transcriptomics. The sampling variability within the tissue had to be accounted for but was not of primary interest. The three mixed effect model methods produced distinct power surfaces in the initial simulation run despite having similar conceptual basis. We attribute these differences to nuances of the software implementation pertaining to calculation of model likelihood and p-values. Still, with all three methods, power depended on the number of patients per group and not ROIs per patient, which is consistent with theoretical considerations [56]. The power of the study depends on the hypothesis/model parameter being tested (in this case, fixed effect regression coefficient for patient group). The number of ROIs per patient affects the ability to accurately estimate the random effects (in this case, patient-specific intercept terms) and not the fixed effect regression coefficients [56]. This represents sharp contrast to the current literature that focuses primarily on spatial variability of expression within the tissue and, accordingly, increasing the number of spots or ROIs per tissue sample [5–7]. Whilst group differences between patients could be addressed with conventional experimental methods such as RNA-sequencing or even qPCR on FACS sorted or laser micro dissected tissue, this presents an impractical barrier in human research when spatial transcriptomics with the NanoString platform enables reliable quantification of gene expression levels from small areas of archived formalin fixed and paraffin embedded tissue.

Due to low amount of input material, there is a risk that some ROIs will fail technical quality control after acquisition of sequencing data. Therefore, we started the simulation with 2 ROIs per patient and endpoint. If one ROI fails technical quality control in a subset of individuals, the remaining replicates should still be sufficient to address the study endpoint. For example, if FLNA expression in 165μm fibrotic niche ROIs was compared between 5 fibrosis progressors and 5 fibrosis-stable individuals (5×2×2 ROI = 20 ROIs) and a third (6) of the ROIs were removed (“failed quality control”) but in a way that did not result in complete loss of the affected patients, then power to detect the group difference with glmer.nb still remained 78.9% based on evaluation of 1,000 synthetic data sets.

glmer.nb yielded similar log2 fold change estimates to the actual spike-in, had good convergence rate and short runtime, so we decided to test primary and secondary endpoints with this method. We set the alpha level to 0.025 (0.05/2 endpoints). We decided that the primary endpoint will be tested first, the secondary endpoint will be tested next and any other analyses such as transcriptome-wide differential expression will be unpowered exploratory. Standard deviations of random intercepts were generally low: median 0.3 (IQR 0.1-0.5) for FLNA in fibrotic niche and median 0.1 (IQR 0.05-0.2) for PON1 in hepatocytes for all N patients and ROI size 165μm in the final simulation study. Type I error in this scenario should be close to the theoretical level [47] and we did not implement any additional steps to control it. However, sensitivity analysis could be conducted via bootstrap method [47]. In addition, expression of summary scores for the lipid metabolism and cytoskeleton markers could be compared between fibrosis progressors, fibrosis regressors and stable fibrosis individuals in sensitivity analysis. We considered this an acceptable approach. If transcriptome-wide significance level was desired, alpha could be set to 1.25e-05, which corresponds to (0.05/2 endpoints)/ca. 2,000 reliably detected genes with 165μm ROI in our reference data. At alpha 1.25e-05, 27 patients per group would be required to achieve 80% power for the primary endpoint (PON1 in hepatocytes) and 15 patients per group would be required for the secondary endpoint (FLNA in fibrotic niche).

### Our sample size calculation is subject to several other considerations

First, the 58 patients with repeated biopsies in Fujiwara *et al*. cohort were at high risk to develop hepatocellular carcinoma and already had detectable liver fibrosis at baseline [23]. Therefore, the hypotheses formulated in this cohort might not generalize to all-comer NAFLD population.

Second, the initial study hypotheses (scores) were based on non-overlapping gene sets. We assumed that the two endpoints pertained to different aspects of biology and different tissue fractions and could therefore be considered independent, which may not necessarily be the case.

Third, NAFLD Brunt fibrosis score reflects the relative amount of fibrotic tissue area within the liver. We assumed that if the amount of fibrotic tissue subjected to sequencing (i.e., the ROI size) is fixed, then we do not need to adjust for fibrosis stage any longer. However, this assumption may not hold true if microscopic structure (e.g., cell composition or cell density) differs within fibrotic niche in patients with different fibrosis severity.

Finally, we focused on the sample size calculation and left other experimental considerations outside of the scope of the current study. For example, identification of ROIs (antibody selection, labelling with fluorescent dyes, reagent concentration etc) and sample randomization (the instrument has throughput of 4-8 slides a day, “GeoMx® Digital Spatial Profiler (DSP) Project Design Guide” document from www.nanostring.com/GeoMxDSP) are separate topics.

In conclusion, NanoString GeoMx is an emerging spatial transcriptomics technology. While some considerations such as definition of regions of interest and their size are technology-specific, we believe that the sample size calculations for spatial transcriptomics, as for any other technology, are predominantly shaped by the biological or clinical study hypothesis. The study hypothesis helps to define the analysis method (statistical model), the associated assumptions and the expected effect size or minimal clinically meaningful difference. In the absence of knowledge-based hypothesis or a dedicated pilot spatial transcriptomics study, the study endpoints can be inferred from prior bulk RNA-sequencing data and re-scaled to tissue fractions corresponding to ROIs in spatial transcriptomics as illustrated in our case study.

## Author contribution

V.A. defined the biological question and guided the study design. M.R. designed and conducted computational analysis and wrote the initial manuscript draft. Both authors revised and approved the final manuscript version.

## Acknowledgments

We would like to thank Magnus Söderberg for valuable discussions on liver histology and Stephanie Ling and Scott Hoffmann for discussions around imaging aspects of data generation.

## Funding

None.

## About the authors

Maria Ryaboshapkina, M.Sc., is a senior scientist at AstraZeneca focusing on applied bioinformatics research in cardiovascular, renal and metabolic diseases.

Vian Azzu, MD PhD, is a principal scientist at AstraZeneca and practicing hepatology clinician in the UK National Health Service. Her research is focused on liver/metabolic diseases and improving patient health outcomes.

## Conflict of interest

The authors are employees of AstraZeneca.

