## Supplementary material for "Sample size calculation for a NanoString GeoMx spatial transcriptomics experiment to study predictors of fibrosis progression in non-alcoholic fatty liver disease": Data S2: 0_calculate_scores.R

```
options(stringsAsFactors = F)

library(Seurat)
library(dplyr) 
library(ggplot2)
library(patchwork)
library(png)
library(viridis)

library(xlsx)

source('functions.R')

markers  =read.xlsx('auxilary_files/markers_updated.xlsx', 1)
markers = markers[, c('cluster', 'gene')]

celltypes = unique(markers$cluster)

gsms = list.dirs(getwd(), recursive = F)
gsms = gsub('.*/', '', gsms)
gsms = gsms[grepl('HC', gsms)]

for(gsm in gsms){

im = Read10X_Image(
  image.dir = getwd(),
  image.name = paste0(gsm, '/spatial/tissue_lowres_image.png'),
  jsonfil = paste0(gsm, '/spatial/scalefactors_json.json'),
  positionfil  = paste0(gsm, '/spatial/tissue_positions_list.csv'),
  filter.matrix = TRUE
)

hu = Seurat::Load10X_Spatial(data.dir = getwd(),
	filename = paste0(gsm, "/filtered_feature_bc_matrix.h5"),
      assay = "Spatial", 
  slice = "slice1", 
  filter.matrix = TRUE, 
  to.upper = FALSE,
  image = im
)

counts = hu@assays$Spatial@counts

hu = SCTransform(hu, assay = "Spatial", verbose = FALSE)
vst = hu@assays$SCT@data
rownames(vst) = toupper(rownames(vst))

scores = matrix(nrow = ncol(vst), ncol = length(celltypes))
rownames(scores) = colnames(vst)
colnames(scores ) = celltypes

for(k in 1:length(celltypes)){
	sym = markers$gene[which(markers$cluster == celltypes[k])]
	subs = as.matrix(vst[which(rownames(vst) %in% sym) ,])
	combined_matrix = rbind(subs,average=apply(subs,2,mean))
	cor_matrix = cor(t(combined_matrix))
	cor_vector = cor_matrix[,dim(cor_matrix)[1]]
	filtered = subs[rownames(subs) %in% names(cor_vector[cor_vector >= 0.1]),]
	scores[,k] = apply(filtered,2,mean)
}

qqECM = quantile(scores[, 'ECM'], 0.85)
qqHep = quantile(scores[, 'Hepatocytes'], 0.15)

ecm_region= rep(0, nrow(scores))
ecm_region[which(scores[, 'Hepatocytes'] <= qqHep &
		scores[, 'ECM'] >= qqECM )] = 1

qqECM = quantile(scores[, 'ECM'], 0.8)
qqHep = quantile(scores[, 'Hepatocytes'], 0.2)

hep_region= rep(0, nrow(scores))
hep_region[which(scores[, 'Hepatocytes'] >= qqHep &
		scores[, 'ECM'] <= qqECM )] = 1


scores = data.frame(barcode = colnames(hu), sample = gsm, hu@meta.data, scores, ecm_region, hep_region)
write.table(scores , paste0('output/', gsm, '_meta.txt'), sep = '\t', row.names = F, quote = F)

}
```
