## Supplementary material for "Sample size calculation for a NanoString GeoMx spatial transcriptomics experiment to study predictors of fibrosis progression in non-alcoholic fatty liver disease": Data S2: 1_check_fibr_area.R

```
gsms = list.dirs(getwd(), recursive = F)
gsms = gsub('.*/', '', gsms)
gsms = gsms[grepl('HC', gsms)]

fils = paste0('output/', gsms , '_meta.txt')

all_meta = read.table(fils[1] , sep = '\t', header = T)

for(i in 2:length(fils)){
meta = read.table(fils[i] , sep = '\t', header = T)
all_meta = rbind(all_meta, meta)
}

tt = table(all_meta$sample , all_meta$hep_region)
print('N hepatocyte spots:')
print(sum(tt[,2]))

qq = apply(tt, 1, function(x) x[2]*100/sum(x))
print('Area occupied by hepatocyte spots:')
print(round(qq, 1))

tt = table(all_meta$sample , all_meta$ecm_region)
print('N fibrotic niche spots:')
print(sum(tt[,2]))

qq = apply(tt, 1, function(x) x[2]*100/sum(x))
print('Area occupied by hepatocyte niche:')
print(round(qq, 1))
```
