## Supplementary material for "Sample size calculation for a NanoString GeoMx spatial transcriptomics experiment to study predictors of fibrosis progression in non-alcoholic fatty liver disease": Data S2: 2_calibration_curve_pt1.R

```
options(stringsAsFactors = F)

library(Seurat)
library(dplyr) 
library(ggplot2)
library(patchwork)
library(png)
library(viridis)

library(xlsx)

source('functions.R')

gsms = c('cHC-1N', 'HCC-1N', 'HCC-2N', 'HCC-3N', 'HCC-4N')
for(gsm in gsms){

	im = Read10X_Image(
  		image.dir = getwd(),
  		image.name = paste0(gsm, '/spatial/tissue_lowres_image.png'),
  		jsonfil = paste0(gsm, '/spatial/scalefactors_json.json'),
 		positionfil  = paste0(gsm, '/spatial/tissue_positions_list.csv'),
  		filter.matrix = TRUE
	)

	hu = Seurat::Load10X_Spatial(data.dir = getwd(),
	filename = paste0(gsm, "/filtered_feature_bc_matrix.h5"),
      	assay = "Spatial", 
  	slice = "slice1", 
  	filter.matrix = TRUE, 
  	to.upper = FALSE,
  	image = im
	)
	counts = hu@assays$Spatial@counts

	pData = read.table( paste0('output/', gsm, '_meta.txt'), sep = '\t', header = T)

	indRow = which(rownames(counts) == 'FLNA')
	indCol = which(colnames(counts) %in% pData[which(pData$ecm_region == 1),1])

	indRow_2 = which(rownames(counts) == 'PON1')
	indCol_2 = which(colnames(counts) %in% pData[which(pData$hep_region == 1),1])

	res = seq(0, 2.9, by = 0.1)
	# added after intital calibration curve run
	res = c(res, 0.43)
	bulk = matrix(nrow = nrow(counts), ncol = length(res))
	colnames(bulk) = paste0(gsm, '_res', res)

	ecm = matrix(nrow = nrow(counts), ncol = length(res))
	colnames(ecm) = paste0(gsm, '_res', res)

	hep = matrix(nrow = nrow(counts), ncol = length(res))
	colnames(hep) = paste0(gsm, '_res', res)

	for(i in 1:length(res)){
		temp = counts
		temp[indRow, indCol] = (median(temp[indRow, indCol])*res[i] + temp[indRow, indCol])
		temp[indRow_2, indCol_2] = (median(temp[indRow_2, indCol_2])*res[i] + temp[indRow_2, indCol_2])
		s = apply(temp, 1, sum)
		bulk[,i] = s
		s = apply(temp[,indCol], 1, sum)
		ecm[,i] = s
		s = apply(temp[,indCol_2], 1, sum)
		hep[,i] = s
	}
	bulk = data.frame(GeneSymbol = rownames(hu), bulk)
	ecm = data.frame(GeneSymbol = rownames(hu), ecm)
	hep = data.frame(GeneSymbol = rownames(hu), hep)

	write.table(bulk, paste0('output/', gsm, '_spiked_in_bulk.txt'), sep = '\t', row.names = F, quote = F)
	write.table(ecm, paste0('output/', gsm, '_spiked_in_ecm.txt'), sep = '\t', row.names = F, quote = F)
	write.table(hep, paste0('output/', gsm, '_spiked_in_hep.txt'), sep = '\t', row.names = F, quote = F)
} 
```
