## Supplementary material for "Sample size calculation for a NanoString GeoMx spatial transcriptomics experiment to study predictors of fibrosis progression in non-alcoholic fatty liver disease": Data S2: 3_calibration_curve_pt2.R

```
options(stringsAsFactors = F)

library(DESeq2)
library(ggplot2)

pats = c('_spiked_in_bulk.txt', '_spiked_in_hep.txt', '_spiked_in_ecm.txt')

for(pat in pats){

gsms = c('cHC-1N', 'HCC-1N', 'HCC-2N', 'HCC-3N', 'HCC-4N')

counts = read.table(paste0('output/', gsms[1], pat), sep = '\t', header = T)

for(i in 2:length(gsms)){
	counts_i = read.table(paste0('output/', gsms[i], pat), sep = '\t', header = T)
	if(nrow(counts) == nrow(counts_i)){
		if(all(counts[,1] == counts_i[,1])){
			counts = cbind(counts, counts_i[, -1])
		}else{print('different format'); stop()}
	}else{print('different format'); stop()}

}

rownames(counts) = counts[,1]
counts = counts[, -1]
counts = round(counts)

meta = data.frame(sample = colnames(counts), patient = gsub('_.*',  '', colnames(counts)), 
res = gsub('.*res', 'res', colnames(counts)))

meta$res = factor(meta$res, levels = paste0('res' , c( seq(0, 0.4, by=0.1), 0.43, seq(0.5, 2.9, by=0.1)) ))
meta$patient = factor(meta$patient)

dds = DESeqDataSetFromMatrix(countData = counts , colData = meta, design = ~ res + patient )
dds = DESeq(dds, minReplicatesForReplace = Inf, fitType = 'mean')

# 0.43 was added after intital calibration curve run

comps = data.frame(control = 'res0', case =paste0('res' , c(0.43, seq(0.1, 2.9, by=0.1))))
ncomps = nrow(comps)

all_deg = matrix(nrow = 0, ncol = 9)

for(i in 1:ncomps){
	case = comps[i, ]$case
	control = comps[i, ]$control

	res = results(dds,contrast=c("res", case , control))
	res = data.frame(GeneSymbol = rownames(res), res)
	res = res[which(res$GeneSymbol %in% c('FLNA' , 'PON1')), ]
	res$contrast = case
	all_deg = rbind(all_deg, res)

}
write.table(all_deg, paste0('output/calibrate_', gsub('_spiked_in_', '_',pat)), sep = '\t' , row.names = F, quote = F)

}
```
