## Supplementary material for "Sample size calculation for a NanoString GeoMx spatial transcriptomics experiment to study predictors of fibrosis progression in non-alcoholic fatty liver disease": Data S2: 4_plot_calibration_curve.R

```
options(stringsAsFactors = F)

library(DESeq2)
library(ggplot2)
library(gridExtra)

ecm_deg = read.table('output/calibrate__ecm.txt', sep = '\t' , header = T)
hep_deg = read.table('output/calibrate__hep.txt', sep = '\t' , header = T)

all_deg = read.table('output/calibrate__bulk.txt', sep = '\t' , header = T)
all_deg = all_deg[which(all_deg$contrast != 'res0.43'), ]
all_deg$contrast = gsub('res', 'r', all_deg$contrast)

p = ggplot(data = all_deg[which(all_deg$GeneSymbol == 'FLNA'), ], aes(x = log2FoldChange, y = contrast )) +
geom_point() +
facet_grid(.~GeneSymbol) +
ylab('Fraction of added counts in fibrotic niche spots') +
xlab('log2FC in bulk liver') +
labs(tag = 'A') +
theme_bw() + theme(panel.grid.minor = element_blank(),
panel.grid.major = element_blank()) +
theme(axis.text = element_text(size = 7, color = 'black'),
axis.title = element_text(size = 7, color = 'black'),
strip.text = element_text(size = 7, color = 'black'),
strip.background = element_blank(),
axis.title.y = element_text(color = 'blue'),
plot.tag = element_text(size = 8, face = 'bold'))

flna_fc = ecm_deg[which(ecm_deg$contrast == 'res2.9' &
ecm_deg$GeneSymbol == 'FLNA'), ]

flna_fc = paste0('log2FC ', round(flna_fc$log2FoldChange, 2), '�' ,
round(flna_fc$lfcSE, 2))

pon1_fc = hep_deg[which(hep_deg$contrast == 'res0.43' &
ecm_deg$GeneSymbol == 'PON1'), ]

pon1_fc = paste0('log2FC ', round(pon1_fc$log2FoldChange, 2), '�' ,
round(pon1_fc$lfcSE, 2))

annot = data.frame(x = 0.385,
 y = c(28, 1), label = c(flna_fc, 'log2FC 0.47'))

p1 = p + geom_vline(xintercept = 0.47, linetype = 'longdash') +
geom_hline(yintercept = 29, linetype = 'longdash', color = 'blue') +
geom_text(data = annot, aes(y = y, x = x, label = label), color = c('blue','black'), size = 2.5) 

p = ggplot(data = all_deg[which(all_deg$GeneSymbol == 'PON1'), ], aes(x = log2FoldChange, y = contrast )) +
geom_point() +
facet_grid(.~GeneSymbol) +
labs(tag = 'B') +
ylab('Fraction of added counts in hepatocyte spots') +
xlab('log2FC in bulk liver') +
theme_bw() + theme(panel.grid.minor = element_blank(),
panel.grid.major = element_blank()) +
theme(axis.text = element_text(size = 7, color = 'black'),
axis.title = element_text(size = 7, color = 'black'),
strip.text = element_text(size = 7, color = 'black'),
strip.background = element_blank(),
axis.title.y = element_text(color = 'blue'),
plot.tag = element_text(size = 8, face = 'bold'))

annot = data.frame(x = 0.57, y = c(5, 1), label = c(pon1_fc, 'log2FC 0.32'))

p2 = p + geom_vline(xintercept = 0.32, linetype = 'longdash')
p2 = p2 + geom_hline(yintercept = 4.3, linetype = 'longdash', color = 'blue') +
geom_text(data = annot, aes(y = y, x = x, label = label), color = c('blue','black'), size = 2.5) 

pdf('output/FigureS2_calibraton_curve.pdf', width = 7.2, height = 3.5)
grid.arrange(p1, p2, ncol = 2)
dev.off()
```
