## Supplementary material for "Sample size calculation for a NanoString GeoMx spatial transcriptomics experiment to study predictors of fibrosis progression in non-alcoholic fatty liver disease": Data S2: 5_simulate_pt1.R

```
options(stringsAsFactors = F)

library(Seurat)
library(dplyr) 
library(ggplot2)
library(patchwork)
library(png)

library(Biobase)
library(reshape2)

library(NanoStringNCTools)
library(GeomxTools)
library(GeoDiff)

library(edgeR)

source('functions.R')

set.seed(900)

gpl = read.table('auxilary_files/GPL32201_GeoMx_Human_Whole_Transcriptome_Atlas.txt', sep = '\t', header = T)

# roi_sizes  = c(2, 4, 9)

# restricted to ROI size for the final simulation
roi_sizes = 9

data(kidney, package = 'GeoDiff')

neg_control = read.table('auxilary_files/negative_controls.txt', sep = '\t', header = T)[,1]

for(roi_size in roi_sizes){

	new_patients = matrix(nrow = 18339, ncol = 0)
	new_patients_meta = matrix(nrow = 0, ncol = 3)

	gsms = c('cHC-1N', 'HCC-1N', 'HCC-2N', 'HCC-3N', 'HCC-4N')
		for(gsm in gsms){
			im = Read10X_Image(
  					image.dir = getwd(),
  					image.name = paste0(gsm, '/spatial/tissue_lowres_image.png'),
  					jsonfil = paste0(gsm, '/spatial/scalefactors_json.json'),
 					positionfil  = paste0(gsm, '/spatial/tissue_positions_list.csv'),
  					filter.matrix = TRUE
				)

			hu = Seurat::Load10X_Spatial(data.dir = getwd(),
				filename = paste0(gsm, "/filtered_feature_bc_matrix.h5"),
      				assay = "Spatial", 
  				slice = "slice1", 
  				filter.matrix = TRUE, 
  				to.upper = FALSE,
  				image = im
				)
			counts = hu@assays$Spatial@counts
			gpl = gpl[which(gpl$ID %in% rownames(counts)), ]
			counts  = counts[match(gpl$ID, rownames(counts)), ]

			pData = read.table( paste0('output/', gsm, '_meta.txt'), sep = '\t', header = T)

			indRow = which(rownames(counts) == 'FLNA')
			indCol = which(colnames(counts) %in% pData[which(pData$ecm_region == 1),1])
	
			counts = counts[, indCol]
			temp = counts
			temp[indRow, ] = counts[indRow, ] + median(counts[indRow, ]) * 2.9
 
			coords = GetTissueCoordinates(hu)
			coords = coords[match(colnames(counts), rownames(coords)), ]
			d = as.matrix(dist(coords))		

			for(iter in 1:250){ 
				for(n_roi in 1:2){ # restricted to params for final simulation
					s = sample(1:nrow(coords), 1)
					nbh = sort(d[s, ])[1:roi_size]
					nbh = names(nbh)
					aggregated_s = rowSums(counts[, nbh])

					ss = sample(1:nrow(coords), 1)
					nbh = sort(d[ss, ])[1:roi_size]
					nbh = names(nbh)
					aggregated_ss = rowSums(temp[, nbh ]) 
					new_patients = cbind(new_patients, aggregated_s , aggregated_ss)

					meta = data.frame(cell = c(paste0('r0_', iter, '_', n_roi), paste0('r29_', iter, '_', n_roi)),
					group = c('res.0', 'res.2.9'),
					patient = c(paste0('r0_',gsm,'_', iter), paste0('r29_',gsm, '_', iter)))

					new_patients_meta = rbind(new_patients_meta, meta)
			}
			} # end making new spots
		} # end iteration over gsm 

	# reformat into object compatible with GeoDiff

	new_patients_meta = data.frame(id = paste0('s', 1:nrow(new_patients_meta)), new_patients_meta) 

	rownames(new_patients_meta) = paste0(new_patients_meta$id, '.dcc')
	colnames(new_patients) = paste0(new_patients_meta$id, '.dcc')
	rownames(new_patients) = rownames(counts)

	fd_example = fData(kidney)	

	fd = data.frame(RTS_ID = rownames(new_patients), TargetName = rownames(new_patients), Module ='RNA-seq', 
			    CodeClass = 'Endogenous', ProbeID  = rownames(new_patients), Negative =FALSE)
	rownames(fd) = rownames(new_patients)

	fd$Negative[which(fd$ProbeID %in% neg_control)] = TRUE
	fd$CodeClass[which(fd$ProbeID %in% neg_control)] = 'Negative'

	pd_example = pData(kidney)

	pd = data.frame(construct = pd_example$construct[1],
				read_pattern =  pd_example$read_pattern[1],
				expected_neg=  pd_example$expected_neg[1],
				patient = new_patients_meta$patient, 
				class = 'ECM', 
				segment = new_patients_meta$id,
				area = pd_example$area[1],
				region = 'ECM',
				group = new_patients_meta$group,
				nuclei = pd_example$nuclei[1])

	rownames(pd) = rownames(new_patients_meta)

	p_example = protocolData(kidney)

	protocol = data.frame(FileVersion = rep( p_example$FileVersion[1], nrow(new_patients_meta)),
			SoftwareVersion = rep(p_example$SoftwareVersion[1], nrow(new_patients_meta)),
			Date = rep(p_example$Date[1], nrow(new_patients_meta)),
			SampleID = rownames(new_patients_meta),
			Plate_ID = rep(p_example$Plate_ID[1], nrow(new_patients_meta)),
			Well = rep(p_example$Well[1], nrow(new_patients_meta)),
			SeqSetId = rep(p_example$SeqSetId[1], nrow(new_patients_meta)),
			Raw = rep(p_example$Raw[1], nrow(new_patients_meta)),
			Trimmed = rep(p_example$Trimmed[1], nrow(new_patients_meta)),          
			Stitched = rep(p_example$Stitched[1], nrow(new_patients_meta)),
			Aligned = rep(p_example$Aligned[1], nrow(new_patients_meta)),
			umiQ30 = rep(p_example$umiQ30[1], nrow(new_patients_meta)),
			rtsQ30 = rep(p_example$rtsQ30[1], nrow(new_patients_meta)),
			DeduplicatedReads = rep(p_example$DeduplicatedReads[1], nrow(new_patients_meta)),
			roi = rep(p_example$roi[1], nrow(new_patients_meta)),
			aoi = rep(p_example$aoi[1], nrow(new_patients_meta)),
			NTC_ID = rep(p_example$NTC_ID[1], nrow(new_patients_meta)),
			NTC  = rep(p_example$NTC[1], nrow(new_patients_meta)) )
	rownames(protocol) = rownames(pd)

	modified = kidney

	e = new.env()
	e$exprs = round(data.matrix(new_patients))

	modified@assayData <- e
	modified@phenoData <- as(pd, 'AnnotatedDataFrame')
	modified@featureData <- as(fd, 'AnnotatedDataFrame')
	modified@protocolData <- as(protocol, 'AnnotatedDataFrame')

	# object for GeoDiff method
	saveRDS(modified, paste0('sim_data/ecm_roi_', roi_size, '.rds'))

	y <- DGEList(counts=round(data.matrix(new_patients)))
	y <- calcNormFactors(y)
	ln.lib.size =effectiveLibSizes(y, log = T)

	FLNA = new_patients[ which(rownames(new_patients) == 'FLNA'),]
	FLNA = unlist(round(FLNA))
	new_patients_meta = data.frame(new_patients_meta, FLNA , ln.lib.size)
	# for excerpt for glmer.nb
	write.table(new_patients_meta, paste0('sim_data/ecm_roi_', roi_size, '.txt'), sep = '\t', row.names = F, quote = F)

}


	
```
