## Supplementary material for "Sample size calculation for a NanoString GeoMx spatial transcriptomics experiment to study predictors of fibrosis progression in non-alcoholic fatty liver disease": Data S2: 7_glmer.nb_ecm.R

```
options(stringsAsFactors = F)

library(lme4)

set.seed(34567)

infils = 'sim_data/ecm_roi_9.txt'

for(infil in infils){
extended = read.table(infil , sep = '\t', header = T)

control = unique(extended$patient[which(extended$group == 'res.0')])
case = unique(extended$patient[which(extended$group == 'res.2.9')])

simres = matrix(nrow = 0, ncol = 10)

n_pats = c(4:20)
iters = c(1:1000)

	comb = matrix(nrow = 0, ncol = 6)

	for(npat in n_pats){
			for(iter in iters){
				newrow = data.frame(fil  = infil , npat, nroi = 2, iter, fc = NA, pv = NA)
				comb = rbind(comb, newrow)
		}
	}

	nro = nrow(comb)

start_time <- Sys.time()

for(k in 1:nro){

	npat = comb$npat[k]

	p = c( sample(control, npat ),
				sample(case, npat ) )

	ind = which(extended$patient %in% p)

	subs = extended [ind,]
	subs$group = factor(subs$group)
	subs$patient = factor(subs$patient)

	tryCatch({
	mo.nb = glmer.nb(FLNA ~ offset(ln.lib.size) + group + (1|patient), data=subs, verbose=FALSE)
	fe = summary(mo.nb)$coefficients

	comb$fc[k] = fe[2, 'Estimate']
	comb$pv[k] = fe[2, 'Pr(>|z|)']
	}, error = function(e){;})
}

outfil = gsub('.txt', '_glmer.nb.txt', infil)
outfil = gsub('sim_data/', 'output/', outfil)
write.table(comb, outfil , sep = '\t', row.names = F, quote = F)

end_time <- Sys.time()
run_time = end_time - start_time
write(run_time, gsub('.txt', '_runtime.txt', outfil))
}
```
