## Supplementary material for "Sample size calculation for a NanoString GeoMx spatial transcriptomics experiment to study predictors of fibrosis progression in non-alcoholic fatty liver disease": Data S2: 9_lrt_ecm.R

```
options(stringsAsFactors = F)

library(GLMMadaptive)

set.seed(34567)

nAGQ = 11

form = as.formula("FLNA ~ offset(ln.lib.size) + group")
form.0 = as.formula("FLNA ~ offset(ln.lib.size)")
formula.random = "~ 1|patient"

infils = 'sim_data/ecm_roi_9.txt'

for(infil in infils){
extended = read.table(infil , sep = '\t', header = T)

control = unique(extended$patient[which(extended$group == 'res.0')])
case = unique(extended$patient[which(extended$group == 'res.2.9')])

simres = matrix(nrow = 0, ncol = 10)

n_pats = c(4:20)
n_rois = 2
iters = c(1:1000)

	comb = matrix(nrow = 0, ncol = 6)

	for(npat in n_pats){
		for(nroi in n_rois){
			for(iter in iters){
				newrow = data.frame(fil  = infil , npat, nroi, iter, fc = NA, pv = NA)
				comb = rbind(comb, newrow)
		}
		}
	}

	nro = nrow(comb)

start_time <- Sys.time()

for(k in 1:nro){

	npat = comb$npat[k]
	nrois = comb$nroi[k]

	p = c( sample(control, npat ),
				sample(case, npat ) )

	ind = which(extended$patient %in% p)


	subs = extended [ind,]
	subs$group = factor(subs$group)
	subs$patient = factor(subs$patient)

	tryCatch({
	model.1 <- mixed_model(form, 
	                       subs, random = as.formula(formula.random), 
                                  family = GLMMadaptive::negative.binomial(), 
                                  nAGQ = nAGQ)

	model.0 <- mixed_model(form.0, 
	                       subs, random = as.formula(formula.random), 
                                  family = GLMMadaptive::negative.binomial(), 
                                  nAGQ = nAGQ)

	fc = fixef(model.1)['groupres.2.9']
	pv = anova(model.0, model.1)$p.value

	comb$fc[k] = fc
	comb$pv[k] = pv

	}, error = function(e){;})
}

outfil = gsub('.txt', '_GLMMadaptive.txt', infil)
outfil = gsub('sim_data/', 'output/', outfil)
write.table(comb, outfil , sep = '\t', row.names = F, quote = F)

end_time <- Sys.time()
run_time = end_time - start_time
write(run_time, gsub('.txt', '_runtime.txt', outfil))

}
```
