## Supplementary material for "Sample size calculation for a NanoString GeoMx spatial transcriptomics experiment to study predictors of fibrosis progression in non-alcoholic fatty liver disease": Data S2: 11_GeoDiff_ecm.R

```
#' ---
#' title: "Main run"
#' author: "Maria Ryaboshapkina"
#' ---

options(warn = -1)
options(message = -1)

library(GeoDiff)
library(dplyr)
library(ggplot2)
library(NanoStringNCTools)
library(GeomxTools)
library(Biobase)
library(reshape2)

set.seed(34567)

rss = 9
fil =  paste0("sim_data/ecm_roi_", rss, ".rds") 

npats_range = c(4:20)
nrois = c(2)
iters = c(1:1000)

	comb = matrix(nrow = 0, ncol = 6)

	for(npat in npats_range){
		for(nroi in nrois){
			for(iter in iters){
				newrow = data.frame(fil, npat, nroi, iter, fc = NA, pv = NA)
				comb = rbind(comb, newrow)
		}
		}
	}

	nro = nrow(comb)

	ecm_all = readRDS(fil)
	control = unique(pData(ecm_all)$patient[which(pData(ecm_all)$group == 'res.0')])
	case = unique(pData(ecm_all)$patient[which(pData(ecm_all)$group == 'res.2.9')])
	pData(ecm_all)$group = factor(pData(ecm_all)$group, levels = c('res.0', 'res.2.9'))

	pData(ecm_all)$patient = factor(pData(ecm_all)$patient)

	ecm_all <- fitPoisBG(ecm_all)
	ecm_all <- fitPoisBG(ecm_all, groupvar = "patient")

	all0probeidx <- which(rowSums(exprs(ecm_all))==0)
	if (length(all0probeidx) > 0) {
    		ecm_all <- ecm_all[-all0probeidx, ]
	}
#	print("call aggreggprobe")
#	ecm_all <- aggreprobe(ecm_all, use = "cor")


	fData(ecm_all)$featureType = "Target"
	fData(ecm_all)$probenum = 1

	ecm_all <- fitNBth(ecm_all)

	pData(ecm_all)$patient = as.character(pData(ecm_all)$patient)

	sum(is.na(fData(ecm_all)$featfact))
	sum(!is.na(fData(ecm_all)$featfact))

	bgMean <- mean(fData(ecm_all)$featfact, na.rm = TRUE)

	r = rowSums(exprs(ecm_all))
	rh = names(sort(r, decreasing = T)[1:1600])

	fh = rownames(ecm_all)[which(rownames(ecm_all) %in% rh &
							fData(ecm_all)$Negative == F)]

start_time <- Sys.time()

 for(sim_run in 1:nro){ 
	tryCatch({

	npat = comb$npat[sim_run]
	nrois = comb$nroi[sim_run]

	p = c( sample(control, npat ),
				sample(case, npat ) )

#	ind = c()
#	for(pp in p){
#		s = pData(ecm_all)$segment[which(pData(ecm_all)$patient == pp)]
#		seg = sample(s, nrois )
#		ii = which(pData(ecm_all)$patient == pp & pData(ecm_all)$segment %in% seg)
#		ind = c(ind, ii)
#	}
	ind = which(pData(ecm_all)$patient %in% p)
	ecm <- ecm_all[, ind]

	pData(ecm)$patient = factor(pData(ecm)$patient)

	ROIs_high <- sampleNames(ecm)

	NBthDEmod = try({
	 	fitNBthDE(form = ~ group,
                       split = F,
                       object = ecm,
			     features_high = fh )
	})

	if(!inherits(NBthDEmod, "try-error")){
	pmat = NBthDEmod$preci1
	}else{pmat = diag(1, 2)}


	NBthmDEmod <- fitNBthmDE(object = ecm,
                         form = ~ group +(1|patient),
                         ROIs_high = ROIs_high,
                         split = FALSE,
                         features_all = c('FLNA', 'LUM'),
                         preci1=pmat,
                         threshold_mean = bgMean,
				 sizefact = ecm$sizefact_fitNBth,
				 sizefact_BG = ecm$sizefact,
                         sizescale = T,
                         controlRandom=list(nu=12, nmh_e=400, thin_e=60))


	coeff <- coefNBth(NBthmDEmod )

	comb$fc[sim_run] = coeff$estimate['groupres.2.9', 'FLNA']
	comb$pv[sim_run] = coeff$p_value['groupres.2.9', 'FLNA']

	}, error = function(e){print(e);} )

} # end loop
	outfil = paste0('output/GeoDiff_', rss, "_ecm.txt")
	write.table(comb, outfil, sep = '\t', row.names = F, quote = F)

end_time <- Sys.time()
run_time = end_time - start_time
write(run_time, gsub('.txt', '_runtime.txt', outfil))
```
