## Supplementary material for "Sample size calculation for a NanoString GeoMx spatial transcriptomics experiment to study predictors of fibrosis progression in non-alcoholic fatty liver disease": Data S2: 13_summarize.R

```
options(stringsAsFactors = F)

fils = c('output/ecm_roi_9_glmer.nb.txt', 'output/ecm_roi_9_GLMMadaptive.txt', 'output/GeoDiff_9_ecm.txt',
'output/hep_roi_9_glmer.nb.txt',
'output/hep_roi_9_GLMMadaptive.txt', 'output/GeoDiff_9_hep.txt')

for(fil in fils){
sim = read.table(fil, sep = '\t', header  = T)
sim$id = paste(sim[,1], sim[,2], sim[,3])

summaryTable = unique(sim[ , 1:3])
summaryTable$id = paste(summaryTable[,1], summaryTable[,2], summaryTable[,3])

summaryTable$no_err_iteration = 0
summaryTable$power_percent = 0
summaryTable$log2FC_median_IQR = 0

for(i in 1:nrow(summaryTable)){
	ind = which(sim$id == summaryTable$id[i])
	temp = sim[ind, ]
	summaryTable$no_err_iteration[i] = sum(!is.na(temp$fc))
	summaryTable$power_percent[i] = sum(temp$pv < 0.025, na.rm = T)*100 / summaryTable$no_err_iteration[i]
	qq = quantile(temp$fc, c(0.25, 0.5, 0.75), na.rm = T)
	qq = round(qq, 2)
	qq = paste0(qq[2], ' (', qq[1], '-', qq[3], ')')
	summaryTable$log2FC_median_IQR[i] = qq
}

summaryTable$flag = 'ok'
summaryTable$flag[which(summaryTable$no_err_iteration < 50)] = 'less than 50 successful iterations'

outn = gsub('.txt', '', fil)
outn = gsub('.*/','', outn )

xlsx::write.xlsx(summaryTable, 'output/simulation_result.xlsx',
	sheetName = outn, append = T, row.names = F)
}
```
