## Supplementary material for "Sample size calculation for a NanoString GeoMx spatial transcriptomics experiment to study predictors of fibrosis progression in non-alcoholic fatty liver disease": Data S2: functions.R

```
Read10X_Image = function (image.dir, image.name = "tissue_lowres_image.png", 
    filter.matrix = TRUE, jsonfil, positionfil) 
{
    image <- readPNG(source = file.path(image.dir, image.name))

    scale.factors = read.csv(jsonfil, header = F)
    nam = gsub(':.*', '', scale.factors[1,])
    nam = gsub('\\}', '', nam)
    nam = gsub('\\{', '', nam) 
    nam = gsub(' ', '', nam) 
    colnames(scale.factors) = nam
    for(i in 1:ncol(scale.factors)){
		x = scale.factors[1,i]
		x = gsub('.*: ', '', x)
    		x = gsub('\\}', '', x)
    		x = gsub('\\{', '', x) 
		x = as.numeric(x)
		scale.factors[1,i] = x
    }

    tissue.positions <- read.csv(file = positionfil, col.names = c("barcodes", 
        "tissue", "row", "col", "imagerow", 
        "imagecol"), header = FALSE, as.is = TRUE, row.names = 1)
    if (filter.matrix) {
        tissue.positions <- tissue.positions[which(x = tissue.positions$tissue == 
            1), , drop = FALSE]
    }

    unnormalized.radius <- as.numeric( scale.factors$fiducial_diameter_fullres ) * 
        as.numeric( scale.factors$tissue_lowres_scalef )

    spot.radius <- unnormalized.radius/max(dim(x = image))

    return(new(Class = "VisiumV1", image = image, 
		scale.factors = scalefactors(spot = as.numeric( scale.factors$tissue_hires_scalef), 
        	fiducial = as.numeric(scale.factors$fiducial_diameter_fullres), 
            hires = as.numeric(scale.factors$tissue_hires_scalef), 
        as.numeric(scale.factors$tissue_lowres_scalef)), 
	  coordinates = tissue.positions, 
        spot.radius = spot.radius))
}
```
