## Supplementary material for "Sample size calculation for a NanoString GeoMx spatial transcriptomics experiment to study predictors of fibrosis progression in non-alcoholic fatty liver disease": Data S2: README.txt

The scripts assume that the reference Visium data sets have been downloaded from PMID 34919432 http://lifeome.net/supp/livercancer-st/data.htm and unzipped into this directory and that two empty directories sim_data/ and output/ have been created.

The folder should look as shown in expected_folder_structure.png

The scripts are provided as an example and rely on existing published software. They are hard-coded for FLNA and PON1 and are not optimized to reduce runtime or RAM usage. RAM usage can be up to ~6 GB for scripts to generate synthetic data sets and GeoDiff. 

Please feel free to modify the scripts to suit your research purpose.
