## Supplementary figures and images for "Sample size calculation for a NanoString GeoMx spatial transcriptomics experiment to study predictors of fibrosis progression in non-alcoholic fatty liver disease"

### expected_folder_structure.PNG

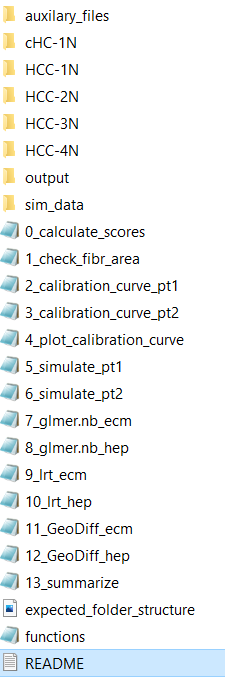
