## Supplementary figures and tables for "Sample size calculation for a NanoString GeoMx spatial transcriptomics experiment to study predictors of fibrosis progression in non-alcoholic fatty liver disease"

**Fig. S1. Correlation between results obtained with univariate linear regression on rlog-transformed counts vs DESeq2 on raw counts in NAFLD GSE135251.** We randomly sampled 28 controls (F0) and 15 cases per more advanced fibrosis stage and evaluated scenarios where log2 fold changes were expected to be small (A), medium (B) and large (C and D). Each data point is a gene.

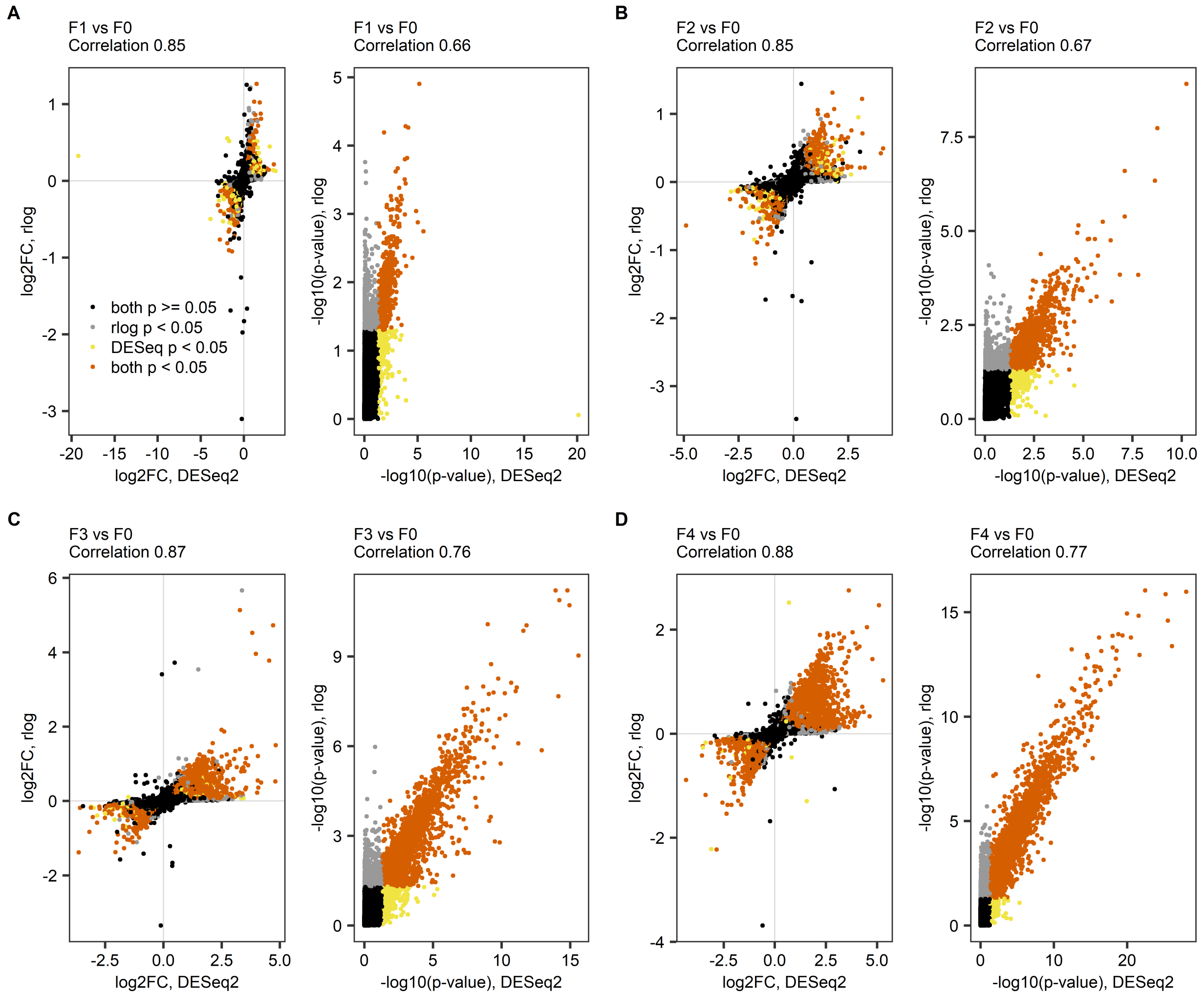

**Fig. S2. Calibration curves.** This analysis was conducted to re-scale expression fold changes from bulk liver to fibrotic niche (A) and hepatocyte fraction (B). The smallest absolute log2 fold change between fibrosis progressors or fibrosis regressors and stable fibrosis individuals in bulk liver was taken to re-scale to the log2 fold change in the corresponding tissue fraction. Numbers on y axis indicate fractions of spiked-in counts. Log2 fold changes were estimated with DESeq2.

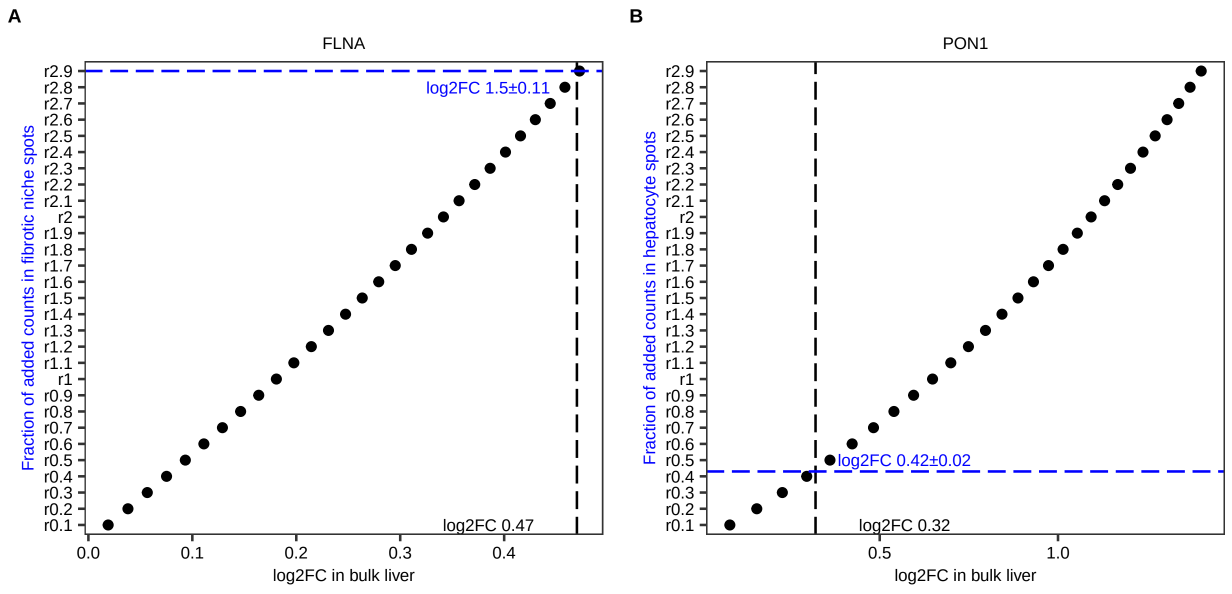

**Fig. S3. Calibration curves in sensitivity analysis.** This analysis was conducted to re-scale expression fold changes from bulk liver to fibrotic niche (A) and hepatocyte fraction (B). The smallest absolute log2 fold change between fibrosis progressors or fibrosis regressors and stable fibrosis individuals in bulk liver was taken to re-scale to the log2 fold change in the corresponding tissue fraction. Numbers on y axis indicate fractions of spiked-in counts. Log2 fold changes were estimated with DESeq2.

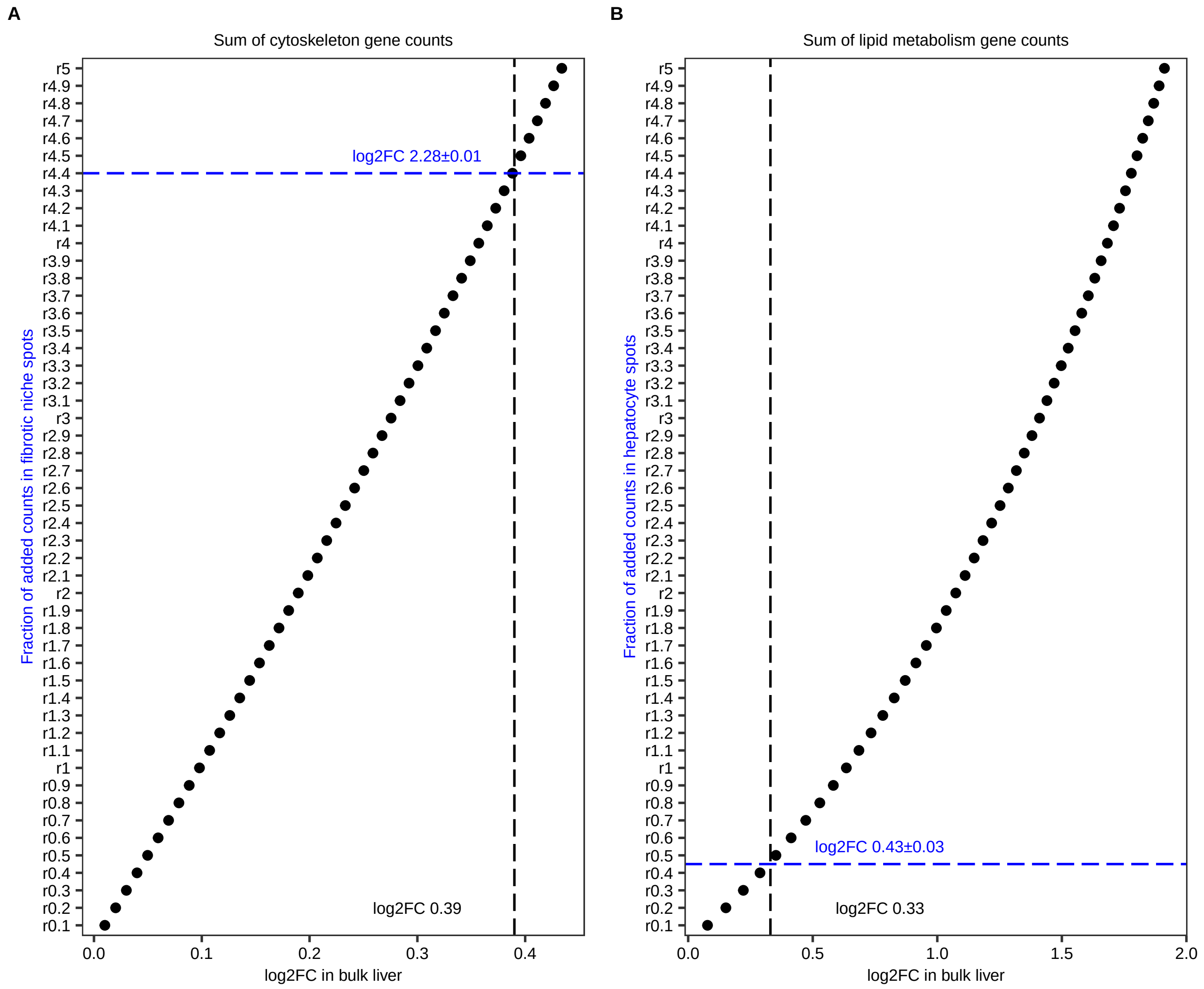

**Table S1.** Results of the sensitivity analysis with glmer.nb, 1,000 iterations per condition and 2 ROIs with 165µm diameter per patient. Number of patients is indicated per group. Alpha = 0.025. Log2 fold change is indicated between spiked-in and unmodified simulated data in (A) fibrotic niche and (B) hepatocyte fraction.

| | A. Sum of cytoskeleton gene counts | | B. Sum of lipid metabolism gene counts | |
|---|---|---|---|---|
| **N patients** | **% Power** | **median (IQR) log2FC** | **% Power** | **median (IQR) log2FC** |
| 4 | 100 | 1.61 (1.45-1.76) | 39.5 | 0.35 (0.19-0.51) |
| 5 | 100 | 1.62 (1.47-1.76) | 42.4 | 0.37 (0.23-0.51) |
| 6 | 100 | 1.61 (1.47-1.75) | 47.8 | 0.37 (0.23-0.5) |
| 7 | 100 | 1.62 (1.49-1.74) | 53 | 0.36 (0.25-0.48) |
| 8 | 100 | 1.61 (1.5-1.74) | 58.4 | 0.36 (0.25-0.47) |
| 9 | 100 | 1.61 (1.5-1.72) | 63.1 | 0.36 (0.26-0.46) |
| 10 | 100 | 1.6 (1.5-1.7) | 65.8 | 0.36 (0.26-0.45) |
| 11 | 100 | 1.62 (1.52-1.72) | 70.3 | 0.36 (0.27-0.45) |
| 12 | 100 | 1.61 (1.53-1.71) | 74.7 | 0.36 (0.27-0.45) |
| 13 | 100 | 1.6 (1.51-1.7) | 76.6 | 0.35 (0.27-0.45) |
| 14 | 100 | 1.62 (1.53-1.69) | 80.8 | 0.36 (0.28-0.44) |
| 15 | 100 | 1.61 (1.53-1.69) | 83.6 | 0.36 (0.28-0.43) |
| 16 | 100 | 1.62 (1.53-1.7) | 84.3 | 0.36 (0.28-0.43) |
| 17 | 100 | 1.61 (1.53-1.69) | 87.6 | 0.36 (0.29-0.43) |
| 18 | 100 | 1.6 (1.54-1.68) | 92 | 0.36 (0.3-0.43) |
| 19 | 100 | 1.61 (1.53-1.68) | 92 | 0.36 (0.29-0.43) |
| 20 | 100 | 1.61 (1.54-1.69) | 94.3 | 0.35 (0.3-0.42) |
